# Genetic overlap between schizophrenia and developmental psychopathology: a longitudinal approach applied to common childhood disorders between age 7 and 15 years

**DOI:** 10.1101/052829

**Authors:** Michel G. Nivard, Suzanne H. Gage, Jouke J. Hottenga, Catherina E.M. van Beijsterveldt, Abdel Abdellaoui, Bart M.L. Baselmans, Lannie Ligthart, Beate St Pourcain, Dorret I. Boomsma, Marcus M. Munafoò, Christel M. Middeldorp

## Abstract

Various non-psychotic psychiatric disorders in childhood and adolescence can precede the onset of schizophrenia, but the nature of this relationship remains unclear. We investigated to what extent the association between schizophrenia and psychiatric disorders in childhood is explained by shared genetic risk factors.

Polygenic risk scores (PRS), reflecting an individual’s genetic risk for schizophrenia, were constructed for participants in two birth cohorts (2,588 children from the Netherlands Twin Register (NTR) and 6,127 from the Avon Longitudinal Study of Parents And Children (ALSPAC)). The associations between schizophrenia PRS and measures of anxiety, depression, attention deficit hyperactivity disorder (ADHD), and oppositional defiant disorder/conduct disorder (ODD/CD) were estimated at age 7, 10, 12/13 and 15 years in the two cohorts. Results were then meta-analyzed, and age-effects and differences in the associations between disorders and PRS were formally tested in a meta-regression.

The schizophrenia PRS was associated with childhood and adolescent psychopathology Where the association was weaker for ODD/CD at age 7. The associations increased with age this increase was steepest for ADHD and ODD/CD. The results are consistent with a common genetic etiology of schizophrenia and developmental psychopathology as well as with a stronger shared genetic etiology between schizophrenia and adolescent onset psychopathology.

A multivariate meta-analysis of multiple and repeated observations enabled to optimally use the longitudinal data across diagnoses in order to provide knowledge on how childhood disorders develop into severe adult psychiatric disorders.

## Introduction

The onset of schizophrenia generally occurs during adolescence or early adulthood^1^, but it is well established that non-psychotic psychiatric symptoms can be present in the period before the first psychotic episode. The prodromal phase is characterized by neurodevelopmental deficits,^2-4^ cognitive learning and memory problems,^5^ and elevated psychiatric symptoms.^6^ Well before the prodromal phase, psychiatric symptoms or disorders are more prevalent in individuals who later develop schizophrenia, as becomes apparent from longitudinal population based cohorts,^7, 8^ retrospective assesments of schizophrenia cases,^9^ and from studies on populations at risk for developing schizophrenia.^10^ Both externalizing symptoms or disorders, including attention deficit hyperactivity disorder, conduct disorder, aggression, and anti-social behavior,^11, 12^ and internalizing symptoms or disorders, including anxiety and depression, are associated with a higher risk of schizophrenia.^7, 8, 11, 13-16^ In sum, these studies indicate that the onset of schizophrenia can be preceded by a broad range of childhood and adolescent psychopathology.

The early detection of schizophrenia can improve outcomes,^8^ and preventive treatment for individuals at risk for schizophrenia can reduce the risk of psychosis.^17, 18^ Insight into the risk factors associated with the predictors of schizophrenia may facilitate early detection. Here, we focused on the role of genetic risk factors. Schizophrenia is highly heritability (approximately 80%^19^) and molecular genetic and twin and family studies generally found evidence for a genetic association between childhood and adult psychopathologies,^20-27^ with one exception.^28^ Consequently, we hypothesized that genetic risk factors for schizophrenia are associated with childhood and adolescent psychopathology. We further expected this association to become stronger from childhood into adolescence, since the the prevalence rates of prodromal symptoms and of psychiatric disorders genetically correlated to schizophrenia (i.e., major depression and bipolar disorder) show a marked increase during adolescence.^1^

We tested these hypotheses with a novel approach which combines the results of multiple polygenic risk score (PRS) analyses of the genetic associations between schizophrenia and longitudinal psychopathology measures into a single multivariate meta-analysis. The PRS were based on the most recent schizophrenia GWA meta-analysis^29^ that yielded 108 genome wide associations, which provides an excellent starting point to investigate the genetic overlap between schizophrenia and other traits (for a review of PRS analyses see ^30-32^). Predictions were tested in two large cohorts with DSM-IV^33^ based measures of anxiety, depression, attention deficit hyperactivity disorder (ADHD) and oppositional deviant disorder and conduct disorder (ODD/CD) assessed at ages 7, 10, 12/13 and 15 years. To simultaneously consider the 192 univariate PRS analyses, a multivariate meta-regression was performed while accounting for the correlations between the multiple polygenic scores and between the psychopathology measures. The multivariate meta-analysis framework provided the opportunity to test differences in the association between schizophrenia PRS and childhood psychopathology between cohorts, disorders and over age.

## Methods

### Subjects

<u>The Netherlands Twin Register (NTR)</u> (www.tweelingenregister.org) follows newborn and adult twins. In the Young NTR (YNTR), twins are registered by their parents and followed from birth onwards. Until age 12, parents complete surveys to report on their twins. From age 14 onwards, information is collected by means of self-report.^34^ In the current study, maternal ratings of childhood psychopathology collected at age 7, 10, and 12 years were analyzed as well as self-report data collected between ages 14-16 years. The number of genotyped children with scores available varied between 1,223 and 2,588 depending on age group (**Supplementary Table 1**). Informed consent was obtained from all participants. The study was approved by the Central Ethics Committee on Research Involving Human Subjects of the VU University Medical Centre, Amsterdam, an Institutional Review Board certified by the U.S. Office of Human Research Protections (IRB number IRB-2991 under Federal-wide Assurance-3703; IRB/institute codes, NTR 03-180).

<u>The Avon Longitudinal Study of Parents And Children (ALSPAC)</u> (www.bristol.ac.uk/alspac/) consists of mothers and their children, born between 1990 and 1991 in the Avon area in southwest England, UK.^35^ The ALSPAC cohort includes maternal ratings of psychopathology at age 7, 10, 13, and 15 and self-ratings at 15 years. The number of genotyped children at each age group varied between 4445 and 6127 (**Supplementary Table 1**). Ethical approval for the study was obtained from the ALSPAC Ethics and Law Committee and the Local Research Ethics Committees. The study website contains details of all data available through a fully searchable data dictionary (www.bris.ac.uk/alspac/researchers/data-access/data-dictionary).

### Measures

In the NTR, psychopathology was measured with DSM-IV based symptom scales^36^ of the age appropriate versions of the Achenbach System of Empirically Based Assessment (ASEBA). For ages 7 to 12, maternal Child Behavior Checklist (CBCL) ratings of the anxiety disorder scale (anxiety), the affective disorder scale (depression), the attention deficit hyperactivity disorder scale (ADHD), and a combined oppositional deviant disorder and conduct disorder ((ODD/CD) scale were analyzed. From age 14 onwards, self-ratings of these scales were analyzed.

In ALSPAC, psychopathology was assessed using the development and wellbeing assessment (DAWBA), which measures the presence of symptoms required for a DSM-IV diagnosis.^37^ Disorders comparable to the ASEBA scales were included in the analyses: any anxiety disorder (anxiety), major depression (depression), attention deficit hyperactivity disorder (ADHD), and combined oppositional deviant disorder and conduct disorder (ODD/CD).^38^ Any anxiety disorder included generalized anxiety disorder, specific phobia, social phobia (at age 7, 10, 13, and 15), separation anxiety disorder (at age 7, 10, and 13), and panic disorder and agoraphobia (at age 15). At ages 7, 10, and 13 all ratings were maternal ratings. At age 15, ADHD and CD/ODD were rated by mothers, and anxiety and depression were self-ratings. The DAWBA yields a diagnosis, but also a more finely grained indicator of disease risk, the DAWBA band. DAWBA band scores, which range from 0 to 5, correspond to probabilities of <0.01%, 0.5%, 3%, 15%, 50%, and >70% of satisfying DSM-IV diagnostic criteria.

### Genotyping

Genotyping and genotype quality control were performed in accordance with common standards to (for a detailed description see **Supplementary Note 1**).

### Polygenic risk scores

Polygenic risk scores (PRS) were calculated by summing the number of risk alleles across all genetic loci (coded as 0,1,2), weighted by the schizophrenia risk conferred by each locus. The risk conferred by each locus was based on the results from the most recent genome-wide association meta-analysis for schizophrenia (PGC-SCZ2, available online:www.med.unc.edu/pgc/downloads).^29^ For all participants, we calculated PRS using LDpred,^39^ a method which accounts for correlations between adjacent genetic loci, and adjusts the risk conferred by each locus accordingly. LDpred further uses a prior expectation for the per locus risk, which is based on the expected degree of polygenicity in a trait. We computed 6 PRS at 6 different priors for the proportion of SNPs with a casual effect (0.01, 0.05, 0.1, 0.25, 0.5, 1). The discovery markers were not pruned and neither were markers a priori eliminated based on thresholds, instead the effect of all markers, the LD between markers and the prior expectation on the degree of polgenicity were leveraged to obtain updated weights for all markers. For each prior an updated set of weights was obtained, which were converted in a polygenic score for each subject. The inclusion criteria for SNPs were minor allele frequency above 5% and high imputation quality (R^2^ > .9). The PRS were scaled to unit variance and mean centered within cohort.

### Statistical analyses

In NTR and in ALSPAC, 96 (4 age bins × 4 disorders × 6 polygenic scores) regression analyses were performed to analyze the prediction of the psychopathology measures by the schizophrenia PRS. Psychopathology measures were scaled to unit variance. As the NTR contained related individuals, the linear regression was performed using a generalized estimation equation with exchangeable background correlations within family, and robust standard errors. This procedure adequately corrects for the presence of related individuals in the sample.^40^ In the ALSPAC sample, an ordered logistic regression was performed since the DAWBA bands are ordered categorical variables. The ordered logistic regression in ALSPAC was transformed to a scale where the underlying latent variable has variance 1. This results in comparable beta’s in the two cohorts, as in both samples an 1 SD increase in the schizophrenia PRS results in an 1 SD increase in the (latent) phenotype., enabling a meta-analysis of regression coefficients from the 96 NTR and 96 ALSPAC analyses. Meta analyses were performed in the metaphor R-package.^41^ In contrast to most meta-analyses, the outcome variables were correlated since within the NTR and ALSPAC the same individuals were repeatedly assessed. The polygenic risk scores are also correlated since they are based on a common set of effect sizes, and only differ in the degree of polygenicity assumed in their construction. These correlations result in dependencies between the parameters to be meta-analyzed. We accounted for this in the meta-analysis by specifying the error covariance matrix as the observed correlations between traits and PRS (see **Supplementary Note 1** for type 1 error simulation and sensitivity analysis).

In the meta-analysis, we tested whether the effect sizes obtained from the 192 univariate PRS analyses departed from zero. We subsequently investigated which variables predicted the strength of the association between psychopathology and schizophrenia PRS in our meta-analysis. In the meta-regression, “age at measurement”, “prior proportion of causal SNPs assumed for PRS”, “cohort” and “disorder” were included as predictors of the strength of the association. Cohort was coded 1 for ALSPAC and 0 for NTR. Age was coded in years over seven (age seven was coded as zero). We considered 4 meta-regression models, including an increasing number of predictors (see Table 1). The most comprehensive model included cohort, age, prior, prior ^2^, isorder, age Xdisorder, age^2^, age^2^Xdisorder. To guard against over fitting, which is a risk in meta-regression, ^42^ we performed 1,000 parametric resamples of the data and performed the model selection on each resample (see **Supplementary Note 1**). We report the proportion of resamples in which each model is selected based on the AIC. We further perform two additional mied effects meta analyses (see **Supplemental Note 1**)

**Table 1:**
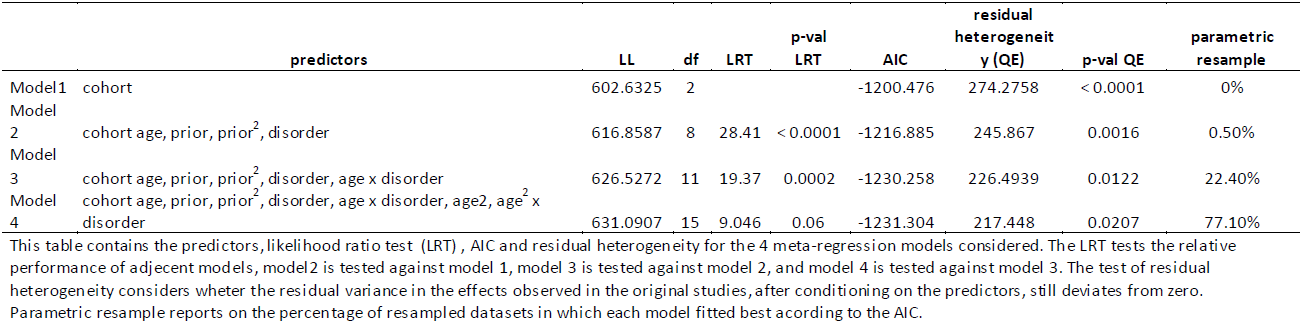
Model fit comparison for themeta-regresslon models.

The regression coefficients obtained in polygenic risk score analyses can be translated to genetic correlations between traits. ^43^ We calculated the genetic correlations between schizophrenia and childhood psychopathology based on the best fitting meta-regression model. Since this transformation relies on assumptions regarding the variance explained by the SNPs and the number of independent SNPs influencing the disorders, we also present the results of the calculations while making different assumptions on the variance explained by all SNPs.

## Results

The descriptives of the psychopathology measures revealed the expected sex differences of adolescent girls scoring on internalizing disorders than boys and boys scoring generally higher than girls on ADHD and ODD/CD (**Supplementary Table 1**-**2** & **Supplementary Note 1**). Consequently, sex was included as a covariate in the PRS analyses.

Model fit statistics, and comparative model fit statistics for the four meta-analytic models are presented in **Table 1**. The model in which the association between childhood psychopathology and schizophrenia PRS are predicted by age, prior, prior^2^, disorder and age × disorder (i.e., a different relationship between disorder and schizophrenia PRS over age) outperforms the basic model which only allows for differences in effect sizes between cohorts. Inclusion of non–linear age effects did not yield an improvement in fit. Likelihood-ratio testing and AIC suggested that Model 3 provided the best balance between parsimony and model complexity. In 77.1% of parametric resamples, model 4 provided the best fit to the data according to the AIC, in 99.5% of the resampled datasets either model 3 or 4 provided the best fit. We proceed to interpret the results of model 3 as the increased in complexity in model 4 by the addition of non-linear age effects yields little extra information. Performing additional effects meta regressions did not substantially change the parameter estimates nor the conclusions drawn based on the meta regression (see **Supplemental Note 1**).

In meta-regression model 3, we tested the degree of polygenicity of the association between schizophrenia and childhood psychopathology by varying the covariate values for prior and prior^2^ while keeping the other covariates fixed at their inverse variance weighted means. The prediction accuracy as a function of prior peaked between the prior values of.50 and 1, suggesting the optimal prior can be found in this range (**Supplementary Figure 1**). This result suggests that the relationship between childhood psychopathology and schizophrenia is highly polygenic in nature, i.e., a large portion of the genome is involved in the relationship between schizophrenia and childhood psychopathology. The forest plot (**Figure 1**), which contains both the empirical and model predicted estimates for the PRS predictions (for PRS prior = 0.50), reveals that the meta-regression predictions are close to the observed PRS regression coefficients.

**Figure 1:**
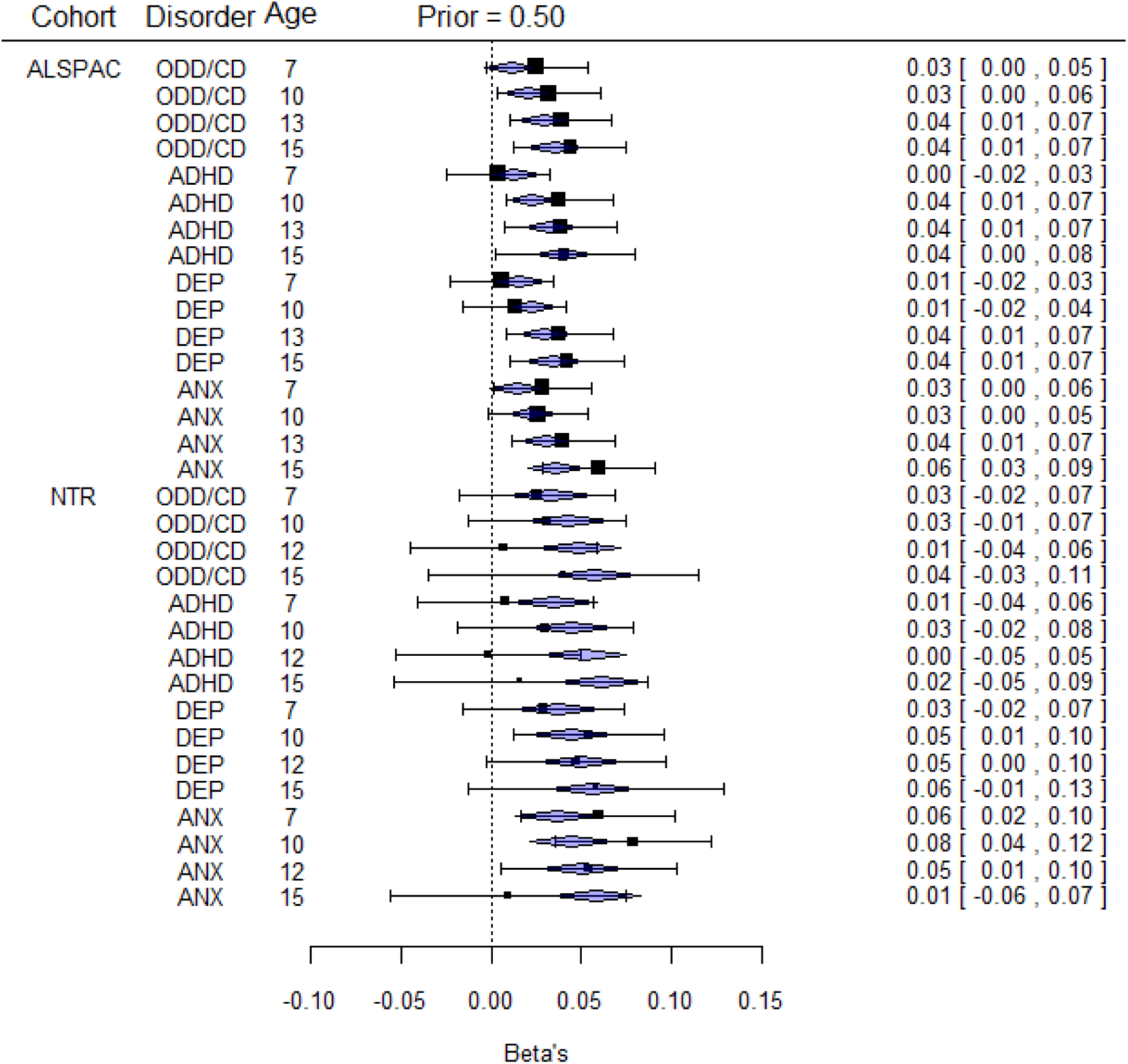
A forest plot of the observed associations between schizophrenia PRS (obtained at prior proportion of causal SNft = 0.05) and of the model predicted associations. The blue polygons indicate the association as predicted from the meta-regression model, while the black square indicates the association as observed in the empirical data. The whiskers indicate the 95% confidence regions around the empirical PRS associations. The resultsare ordered byincreasing age foreach disorder, with in the top halve the results in the ALSPAC cohort and in the bottom halve the results for the NTR cohort.

**Figure 2** shows, based on model 3, the associations between schizophrenia PRS and childhood psychopathology as a function of age, and age × disorder, while keeping all other predictors (i.e, cohort, prior and prior^2^) fixed at their respective inverse variance weighed mean value. The associations increases with age, confirming our hypothesis that the genetic relationship between schizophrenia and developmental psychopathology is stronger in adolescence than in early childhood. The parameter estimates in model 3 (**Table 2**) further indicate that the association with schizophrenia at age 7 was highest for depression (0.0262, Z= 2.227, p < 0.03) and significantly lower for ODD/CD compared to depression (Z=-2.49., p < 0.02). The predictions for ADHD (Z = 1.61, P < 0.11) and anxiety (Z = - .38, p = .70) did not differ with depression. The increase in association with schizophrenia with age was significant for depression (Z = 2.93, p < 0.003) and stronger for ADHD (Z = 4.18, p < 0.001) and ODD/CD (Z = 2.17, p < 0.03) but not anxiety compared to depression (Z = 0.97, p = .30).Further sensitivity analyses showed that the model selection and model parameters were robust to considerable misspecification of the error covariance matrix (see **Supplementary Note 1**).

**Figure 2:**
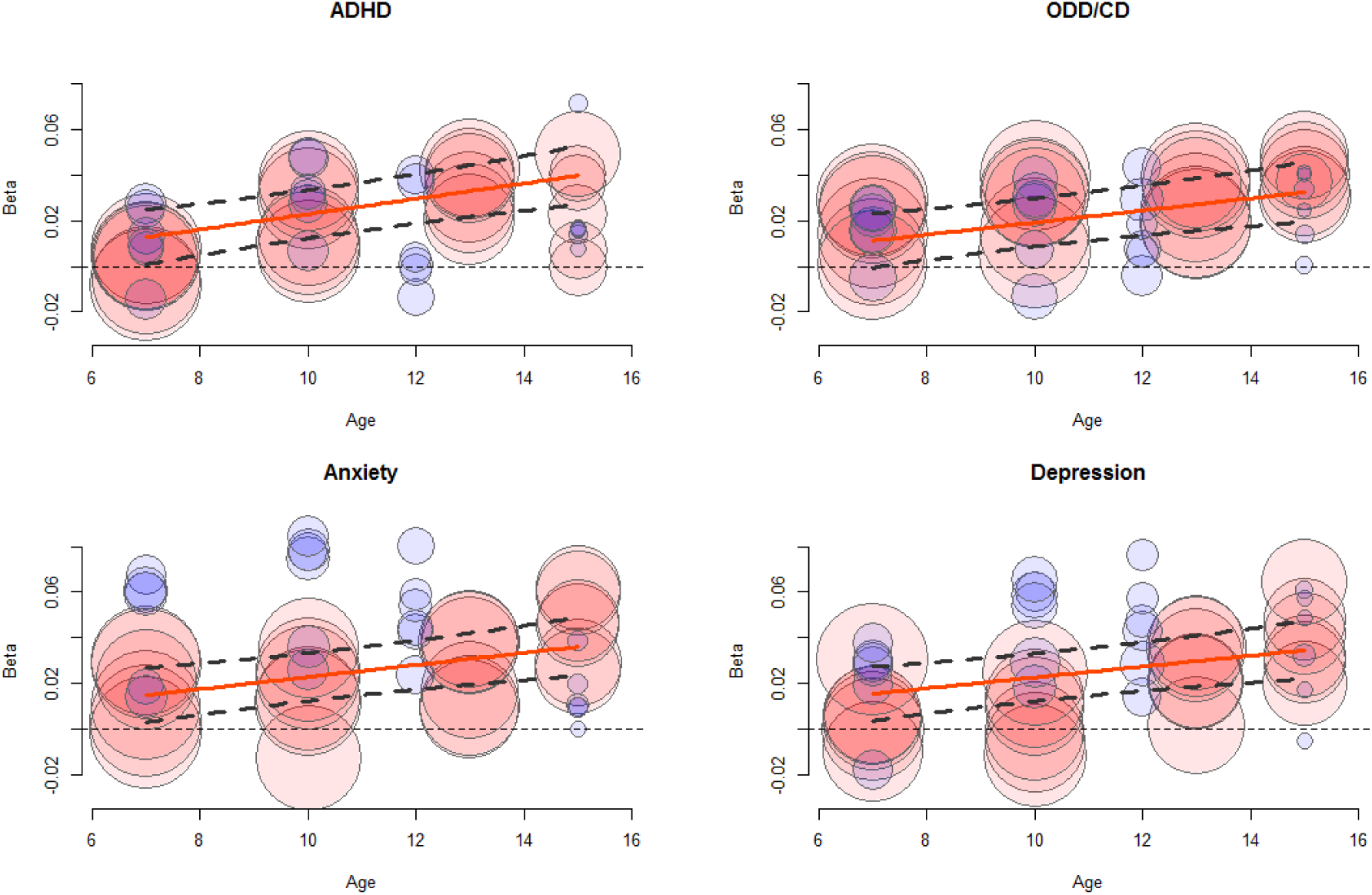
Bubble plot showing the effect of age on the association between schizophrenia PRS and childhood psychopathology, split per disorder. Ordes indicate the observed effect sizes in the univariate regression analyses (ALSPAC in blue, NTR in red). The size of the circles is proportional to the inverse of the variance, and thus larger circles reflect more accurate estimates. The solid line reflects the meta-regression fitted effect size and the dashed lines indicate the upper and lower 95% confidence interval around the meta-regression line.

**Table 2:**
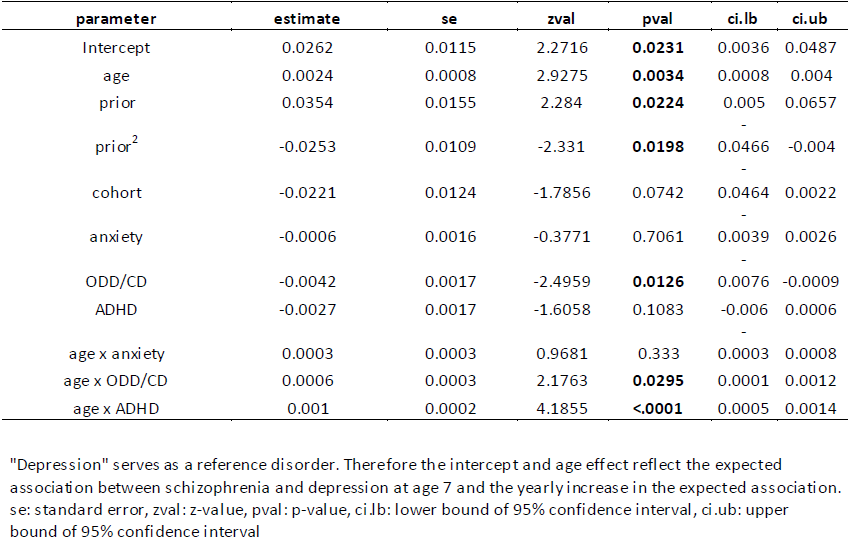
Parameter estimates and test statistics for nieta-regression model 3.

Finally, based on the relationship between the outcomes of PRS analyses and genetic correlations as described by Dudbridge,^43^ we computed the expected genetic correlation between developmental psychopathology and schizophrenia as a function of age and split over disorders based on the beta’s obtained in the meta-regression (for details see: **Supplementary Note 1**). We assumed that 15% of the variance in childhood psychopathology is captured by the genetic markers used to compute the scores,^23, 44, 45^ that 35% of variance in the schizophrenia liability is explained by the markers included in the score, ^24, 46^ and that 200,000 independent genetic effects are captured by the markers included in the PRS. Given these assumptions genetic correlations increase from around 0.10 at age 7 and around 0.25 at age 16, differences in genetic correlations with schizophrenia between disorders were modest (**Figure 3**). The influence of assuming lower (10%) or higher (20%) true variance explained by all measured genetic markers in childhood psychopathology on the estimated genetic correlation is quantified in **Supplementary Figures 2**-**3**.

**Figure 3:**
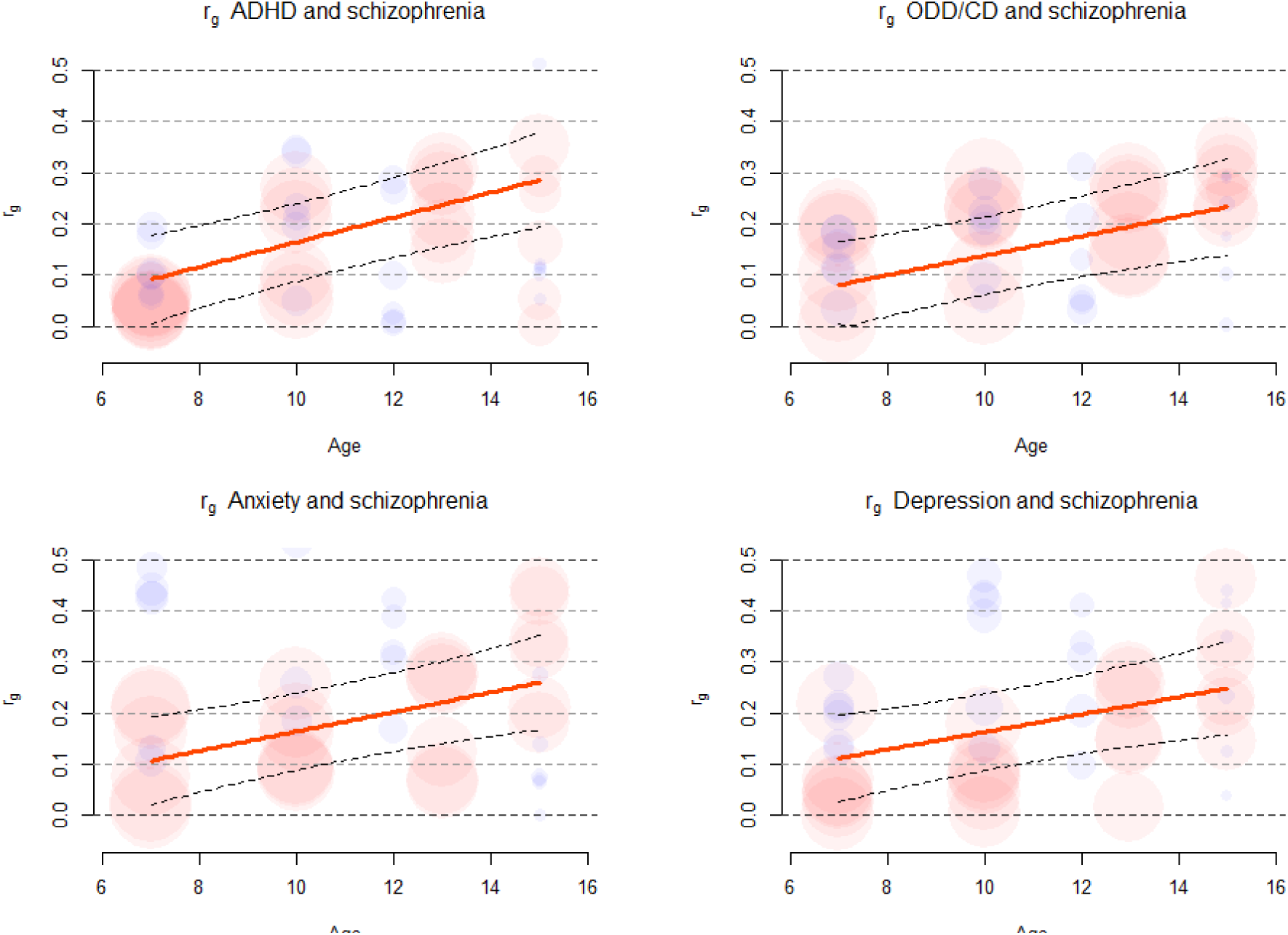
Bubble plot of the approximated genetic correlations between schizophrenia and childhood psychopathology per disorder. We assume the variance in childhood psychopathology explained by all markers used to construct the PRS is 15% and constant over disorders and age. We further assume the PRS captures 200.000 independent genetic effects. Grdes indicate the transformed observed regression coefficients to genetic correlations (ALSPACin blue, NTR in red). The size of the tirdes is proportional to the inverse of the variance, and thus larger cirdes reflect more accurate estimates. The solid line reflects the genetic correlation and the dashed lines indicate the uppe rand lower 95% confidence interval around the genetic correlation, quantifyng the uncertainty in the meta-regression but not in the variance in childhood psychopathology explained by all measured markers, or the estimate of the number of independent markers.

## Discussion

We investigated whether associations between schizophrenia PRS and childhood psychopathology were explained by shared genetic risk factors. For childhood and adolescent anxiety, depression, ADHD and OCC/CD we found the association to be significant across all ages. The genetic overlap between schizophrenia PRS and childhood psychopathology became stronger with age. The associations differed between disorders, with e.g. a weaker association for ODD/CD at age 7, and a stronger age related increase for ADHD. Given reasonable assumptions, the observed polygenic risk predictions translate to genetic correlations between schizophrenia and psychopathology increasing from ∼0.1 in childhood to ∼.25 in adolescence. We further found evidence for a high degree of polygenicity in the relationship between schizophrenia PRS and childhood psychopathology, as was evident from the increase in effect with an increase in the polygenicity prior used in computing the polygenic risk scores.

Strengths of our study were the substantial sample sizes of the discovery and target samples. The schizophrenia PRS were based on a large discovery set, a GWAS which revealed 108 genome wide significant loci.^29^ The target samples varied between 5,354 and 8,253 at different ages, which is substantially higher than the required number of ∼2,000 subjects generally indicated as sufficient for PRS analysis.^30^ Innovative strengths of our analyses were the explicit modeling of all univariate analysis results in the meta-regression approach which accounted for the covariance between disorders and ages, correction for cohort specific effects and the simultaneous consideration of multiple risk scores trained at different priors.

The use of different measures of psychopathology in ALSPAC and NTR could be considered either a limitation or strength. Both measures were based on maternal and self-ratings that are consistently related to clinical DSM-IV diagnoses.^36, 37, 47^ As we observed a significant prediction in both cohorts, results can be generalized across different indices of childhood psychopathology. Several genetic studies on ADHD have also shown that genetic factors for clinically diagnosed ADHD overlap with genetic factors influencing continuous ADHD measures in the general population.^48-50^ These results indicate that combining clinical and population based data in a single study can also be a way to increase sample size and thus statistical power. Another limitation that concerns longitudinal studies is the dropout over the years. We analyzed whether the schizophrenia PRS predicted non-participation and observed significant associations between schizophrenia PRS and non-participation at age 15 in both cohorts and at all ages in ALSPAC (**Supplementary Table 3**). We further observed that non-participation was related to a higher score on psychopathology scales at an earlier age in ALSPAC; this especially holds for externalizing and ADHD at age 13 and 15. In NTR non-participation at age 15 was related to externalizing and depression at age 12 (**Supplementary Table 4**). As those with higher PRS and higher psychopathology scores at earlier ages are more likely to dropout, we expect that the dropout introduces downward bias in the estimated relationship between schizophrenia and childhood psychopathology. Note that only longitudinal analyses can provide insight into the influence of dropout on the estimate genetic relationship between traits, while in univariate studies a failure to participate results in the absence of genetic data and thus the influence of failure to participate cannot be quantified. For more comprehensive genetically informed dropout analysis of the ALSPAC data see:^51^.

Previous research focusing on the genetic overlap between schizophrenia and (childhood) psychopathology is largely in line with our results.^24, 25, 27, 28, 52^ Three differences are noteworthy. Another study in the ALSPAC sample focused on psychiatric symptoms at age 15 and found that schizophrenia PRS predicted anxiety disorder and negative symptoms, but not depressive disorder and psychotic experiences. The difference with the current results for depression may be explained by improvements in the method used to compute the polygenic scores and in the definition of the phenotype.^27^ Two studies by the Psychiatric Genomics Consortium Cross Disorder Group detected strong correlations between major depressive disorder, bipolar disorder and schizophrenia, but no genetic correlation between ADHD and schizophrenia. ^24, 25, 52^ The latter is probably explained by the smaller sample size at the time, as is confirmed by recent analyses which detect a significant association between ADHD and schizophrenia.^52^ Finally, a study^28^ analyzing attention problems and impulsivity, anxiety and a general tendency for psychopathology yielded no significant associations. Yet, the direction of the effects was positive (i.e., higher schizophrenia risk predicted higher scores). As that study comprised polygenic risk scores for 13 traits as well as 50 outcome variables, the multiple-testing burden was considerably higher than in the current study resulting in lower power to detect an effect. Overall, the evidence suggests the presence of a positive genetic correlation between schizophrenia and psychopathology in childhood and in adulthood. Stronger associations are generally reported in adulthood than in childhood. Our findings add a longitudinal and multivariate perspective, and provide evidence for strong polygenicity in the relationship between schizophrenia and childhood psychopathology.

While it is tempting to discuss significant prediction of childhood and adolescent psychopathology by schizophrenia polygenic risk in terms of future clinical risk prediction, we instead offer a word of caution. Genetic prediction is in its infancy. The predictions made, as of yet, are very weak predictors of individual clinical outcomes. The absolute upper bound for genetic risk prediction is easily found by considering the disease concordance for monozygotic, i.e., genetically nearly identical, twins, which is ∼50% for schizophrenia. The concordance is a function of shared genetic and environmental exposures both pre and post-natal.^53, 54^ Genetic clinical risk prediction will improve, but will not exceed this upper bound.

The current study shows how longitudinal analysis can provide insight into how genetic factors exert their effects across diagnostic boundaries and over ages. By explicitly testing sources of heterogeneity (age and disorder) we shed light on the effects of schizophrenia risk genes during childhood. Our findings suggest that there are sets of SNPs broadly influencing psychopathology across ages in the general population, and that there are sets of SNPs of which the effect is either limited puberty or increases in puberty. Our results indicate that age-sensitive genome-wide meta-analysis of repeated measures, in either case-control or population based samples could well identify genetic variants. Some of these genetic variants will increase an individual’s vulnerability for psychopathology and may be associated with persistence of symptoms from childhood into adolescence and adulthood, while other variants can be identified that have an age or disorder dependent effect on psychopathology. Identifying not only which variants influence psychopathology but also at what age can aid to focus translational studies on developmental processes.

To conclude, our study shows how genetic risk factors for schizophrenia are of increasing importance during childhood and adolescence and demonstrate the value of longitudinal studies across diagnostic boundaries to increase our insight into the etiology of severe psychiatric disorders.

## Acknowledgments.

We are extremely grateful to all the families who took part in this study, the midwives for their help in recruiting them, and the whole ALSPAC team, which includes interviewers, computer and laboratory technicians, clerical workers, research scientists, volunteers, managers, receptionists and nurses. The UK Medical Research Council and the Wellcome Trust (Grant ref: 102215/2/13/2) and the University of Bristol provide core support for ALSPAC. This work was supported in part by the Medical Research Council Integrative Epidemiology Unit at the University of Bristol (MC_UU_12013/6). This publication is the work of the authors and Michel G. Nivard will serve as guarantor for the contents of this paper. ALSPAC GWAS data was generated by Sample Logistics and Genotyping Facilities at the Wellcome Trust Sanger Institute and LabCorp (Laboratory Corporation of America) using support from 23andMe. We are equally grateful to the NTR twins, their parents and other family members who participate in the NTR research. The NTR gratefully acknowledges support from “Genetic influences on stability and change in psychopathology from childhood to young adulthood”(ZonMW 912-10-020) and “Genetics of Mental illness” (ERC-230374). MGN is supported by Royal Netherlands Academy of Science Professor Award (PAH/6635). The Netherlands Organization for Scientific Research (NWO) and MagW/ZonMW grants Middelgroot-911-09-032, Center for Medical Systems Biology (CSMB, NWO Genomics), NBIC/BioAssist/RK(2008.024), Biobanking and Biomolecular Resources Research Infrastructure (BBMRI–NL, 184.021.007), VU University’s Institute for Health and Care Research (EMGO+) and Neuroscience Campus Amsterdam (NCA). Part of the genotyping and analyses were funded by the Genetic Association Information Network (GAIN) of the Foundation for the National Institutes of Health, Rutgers University Cell and DNA Repository (NIMH U24 MH068457-06), the Avera Institute, Sioux Falls, South Dakota (USA) and the National Institutes of Health (NIH R01 HD042157-01A1, MH081802, Grand Opportunity grants 1RC2 MH089951 and 1RC2 MH089995).

## Conflicts of interest

The authors declare no conflict of interest.

Supplementary information is available at Molecular Psychiatry’s website

## Author contributions

MGN & CMM devised the study design

CMM, MR M & DIB supervised the research

MGN, SHG performed the analysis

MGN,CMM, DIB, MR M, SHG wrote the manuscript

AA, BSP, CEMB, JJH, BMLB, LL contributed to the analyses, genotypes and phenotypes.

---

**Supplementary Figure 1:**
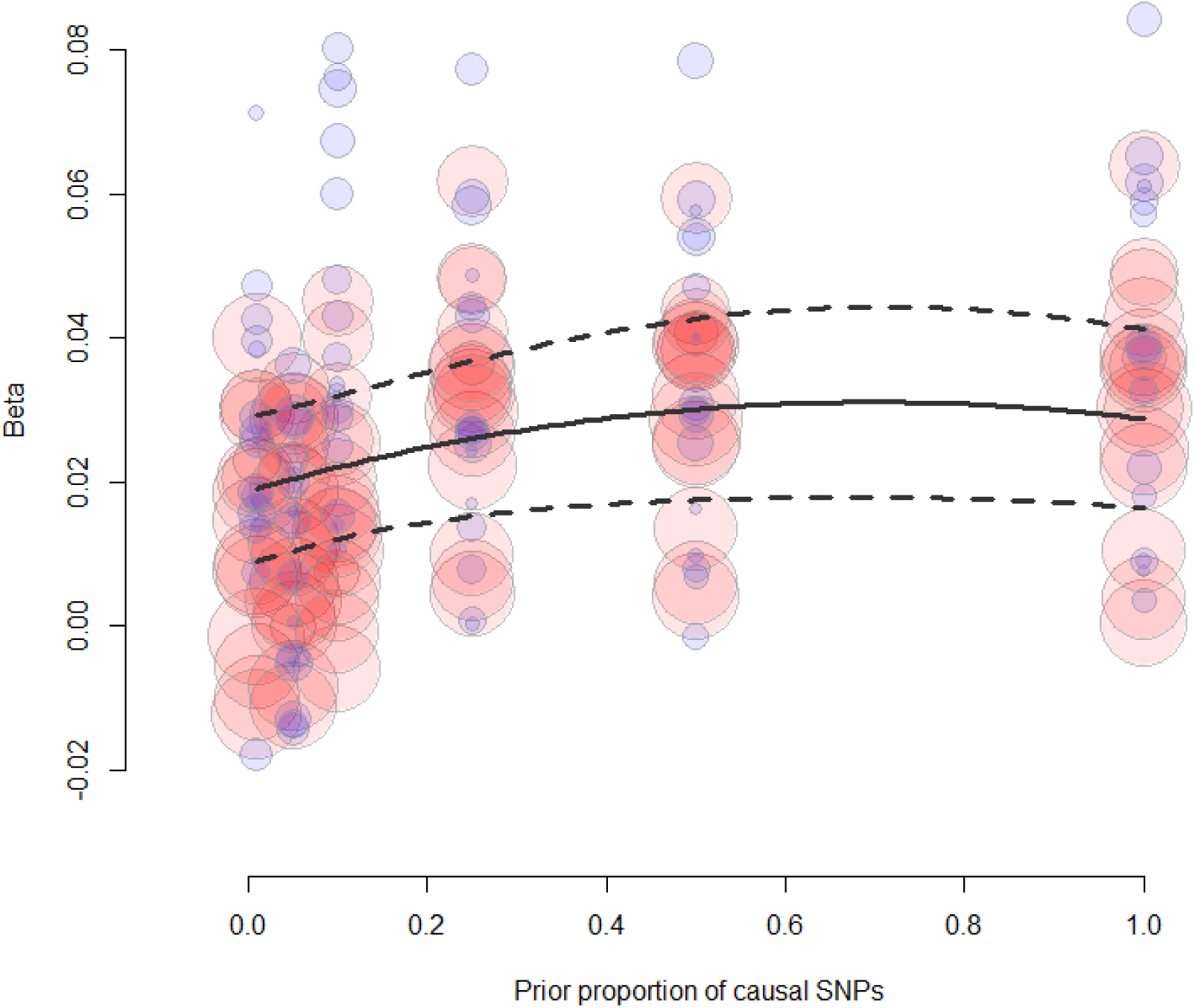
Bubble plot of the relationship between the polygenidty prior and the effectsize in the polygenic risk score analyses. Not the increase in effect size with the increase in polygenicity prior. The solid line indicates the best fit obtained from the meta-regression model (model 3) where the dashed lines reflect the upper and lower confidence bounds.

**Supplementary Figure 2.**
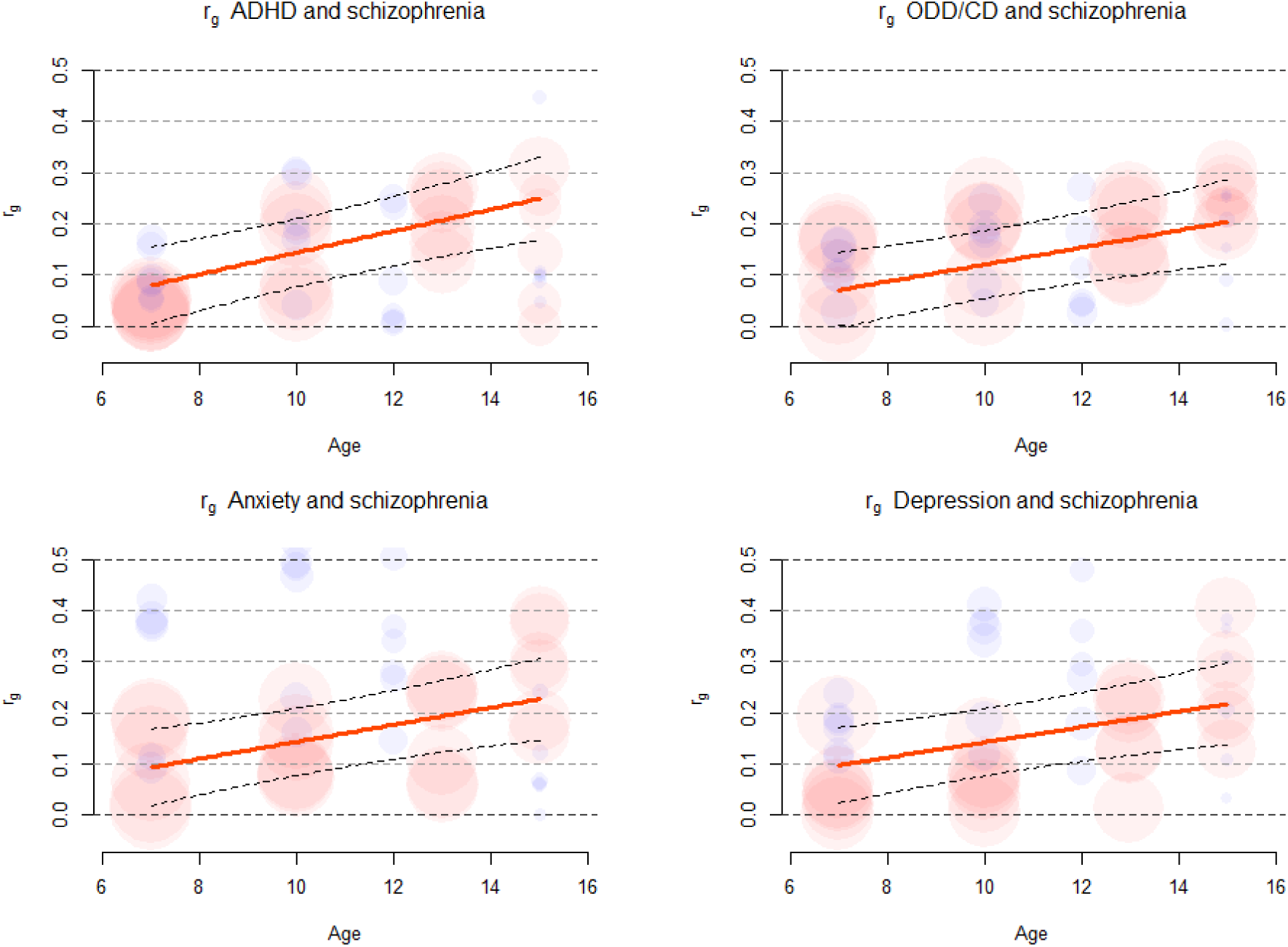
Bubble plot of the approximated genetic correlations between schizophrenia and childhood psychopathology per disorder given the assumptions described in Supplementary Note 1. In this figure we assume the variance explained by all markers in childhood pscyhopathology is constant and 20%. Circles indicate the transformed observed regression coefficients to genetic correlations (ALSPAC in blue, NTR in red). The size of the circles is proportional to the inverse of the variance, and thus larger Circles reflect more accurate estimates. The solid line reflects the genetic correlation and the dashed lines indicate the upper and lower 95% confidence interval around the genetic correlation, quantifying the uncertainty in the meta-regression but not in the variance in childhood psychopathology explained by all measured markers, or the estimate of the number of independent markers.

**Supplementary Figure 3.**
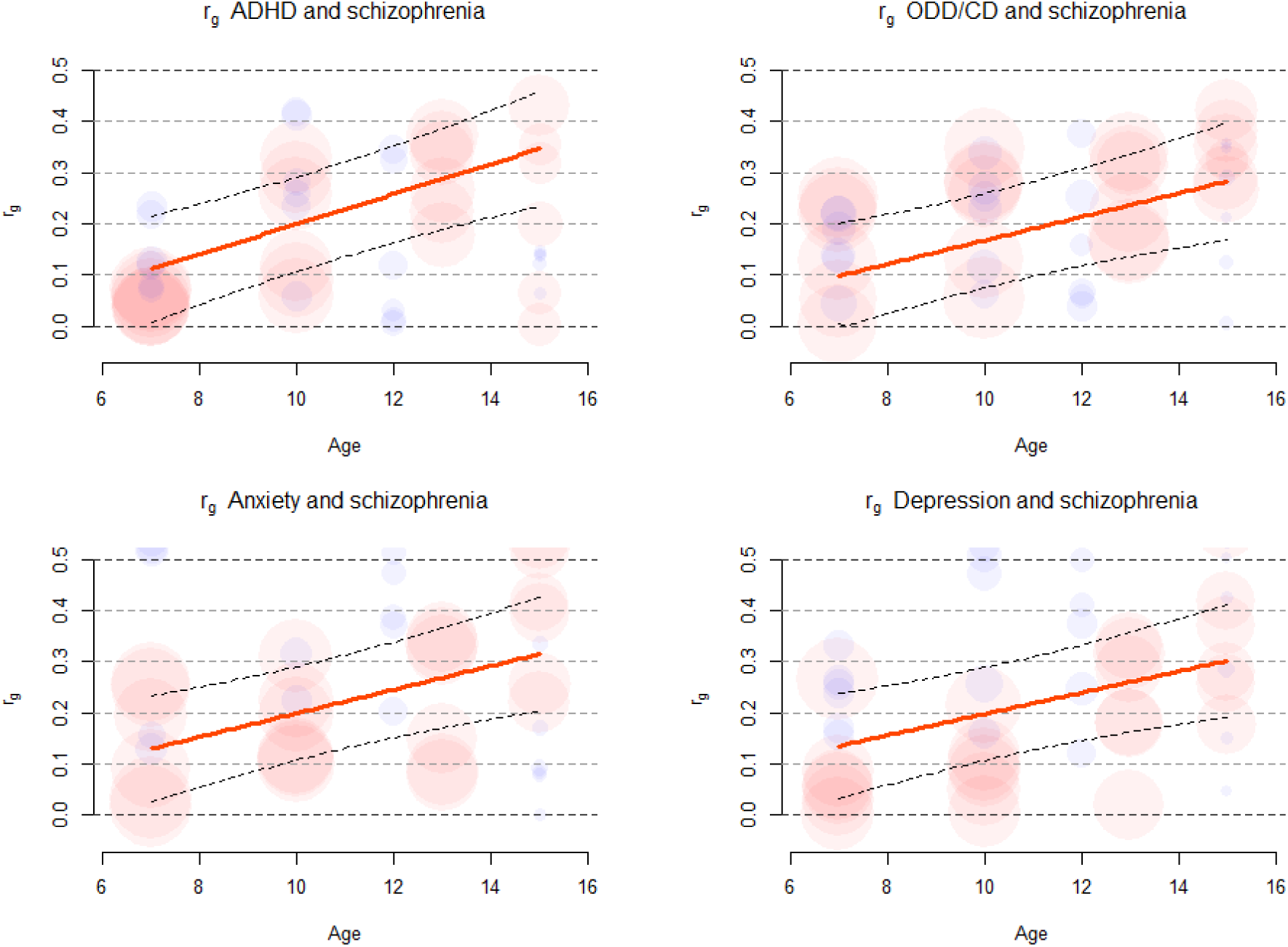
Bubble plot of the approximated genetic correlations between schizophrenia and childhood psychopathology per disorder given the assumptions described in Supplementary Note 1. In this figure we assume the variance explained by all markers in childhood pscyhopathology is constant and 10%. Circles indicate the transformed observed regression coefficients to genetic correlations (ALSPACin blue.NTRin red). The size of the circles is proportional to the inverse of the variance, and thus larger Circles reflect more accurate estimates. The solid line reflects the genetic correlation and the dashed lines indicate the upper and lower 95% confidence interval around the genetic correlation, quantifying the uncertainty in the meta-regression but not in the variance in childhood psychopathology explained by all measured markers, or the estimate of the number of independent markers.

**Supplementary Table 1:** Sample sizes per age group for the NTR, ALSPAC and combined

| **Age** | **NTR** | **ALSPAC** | **Total** |
| --- | --- | --- | --- |
| Age 7 | 2588 | 6127 | 8715 |
| Age 10 | 2487 | 5934 | 8421 |
| Age 12 | 2314 | 5496 | 7810 |
| Age 15 | 1223 | 4445 | 5668 |

**Supplementary Table 2:**
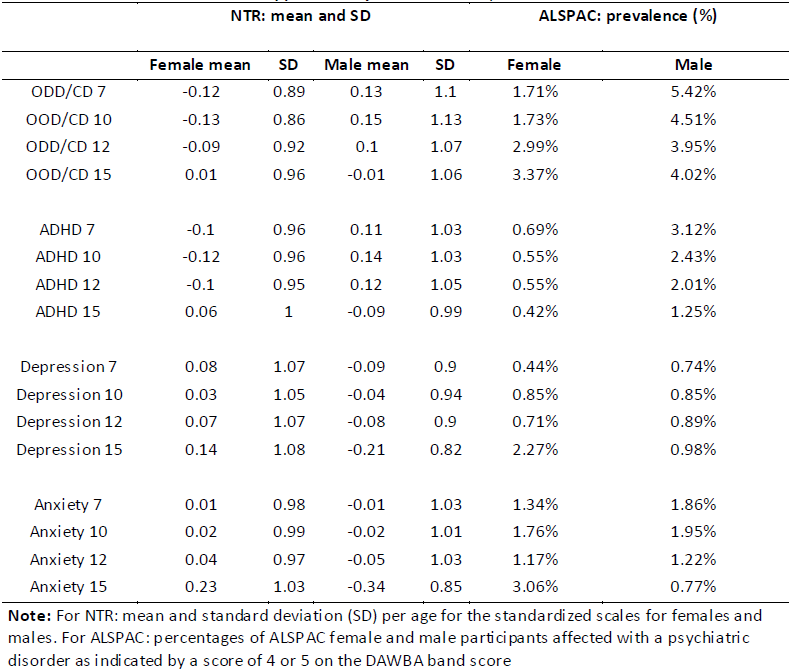
Descriptives

|  | NTR: mean and SD |  |  |  | ALSPAC: prevalence (%) |  |
| --- | --- | --- | --- | --- | --- | --- |
|  | Female mean | SD | Male mean | SD | Female | Male |
| ODD/CD 7 | -0.12 | 0.89 | 0.13 | 1.1 | 1.71% | 5.42% |
| OOD/CD 10 | -0.13 | 0.86 | 0.15 | 1.13 | 1.73% | 4.51% |
| ODD/CD 12 | -0.09 | 0.92 | 0.1 | 1.07 | 2.99% | 3.95% |
| OOD/CD 15 | 0.01 | 0.96 | -0.01 | 1.06 | 3.37% | 4.02% |
| ADHD 7 | -0.1 | 0.96 | 0.11 | 1.03 | 0.69% | 3.12% |
| ADHD 10 | -0.12 | 0.96 | 0.14 | 1.03 | 0.55% | 2.43% |
| ADHD 12 | -0.1 | 0.95 | 0.12 | 1.05 | 0.55% | 2.01% |
| ADHD 15 | 0.06 | 1 | -0.09 | 0.99 | 0.42% | 1.25% |
| Depression 7 | 0.08 | 1.07 | -0.09 | 0.9 | 0.44% | 0.74% |
| Depression 10 | 0.03 | 1.05 | -0.04 | 0.94 | 0.85% | 0.85% |
| Depression 12 | 0.07 | 1.07 | -0.08 | 0.9 | 0.71% | 0.89% |
| Depression 15 | 0.14 | 1.08 | -0.21 | 0.82 | 2.27% | 0.98% |
| Anxiety 7 | 0.01 | 0.98 | -0.01 | 1.03 | 1.34% | 1.86% |
| Anxiety 10 | 0.02 | 0.99 | -0.02 | 1.01 | 1.76% | 1.95% |
| Anxiety 12 | 0.04 | 0.97 | -0.05 | 1.03 | 1.17% | 1.22% |
| Anxiety 15 | 0.23 | 1.03 | -0.34 | 0.85 | 3.06% | 0.77% |

**Note:** For NTR: mean and standard deviation (SD) per age for the standardized scales for females and males. For ALSPAC: percentages of ALSPAC female and male participants affected with a psychiatric disorder as indicated by a score of 4 or 5 on the DAWBA band score

**Supplementary Table 3:**
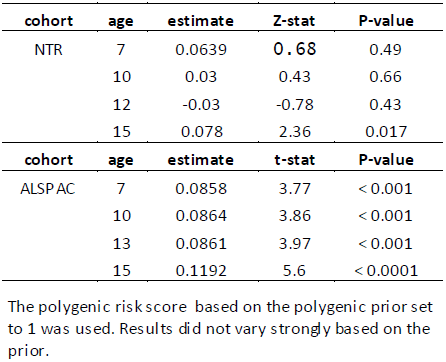
Prediction of non-participation based on the schizophrenian PRS.

**Supplementary Table 4:** Prediction of non-participation based on psychopathology scores at an earlier time point.

| cohort | Outcome | Predictor | estimate | Z-stat | P-value |
| --- | --- | --- | --- | --- | --- |
| NTR | missingness | ODD/CD |  |  |  |
|  | age 10 | age 7 | -0.021 | -1.52 | 0.13 |
|  | age 12 | age 10 | -0.002 | -0.09 | 0.93 |
|  | age 15 | age 12 | 0.034 | 2.03 | 0.043 |
| NTR | missingness | ADHD |  |  |  |
|  | age 10 | age 7 | -0.002 | -0.46 | 0.65 |
|  | age 12 | age 10 | -0.002 | -0.11 | 0.91 |
|  | age 15 | age 12 | 0.012 | 1.05 | 0.29 |
| NTR | missingness | Depression |  |  |  |
|  | age 10 | age 7 | -0.014 | -1.77 | 0.077 |
|  | age 12 | age 10 | -0.02 | -1.06 | 0.29 |
|  | age 15 | age 12 | 0.041 | 2.61 | 0.009 |
| NTR | missingness | Anxiety |  |  |  |
|  | age 10 | age 7 | -0.0089 | -0.92 | 0.36 |
|  | age 12 | age 10 | -0.005 | -0.5 | 0.62 |
|  | age 15 | age 12 | 0.016 | 1.04 | 0.3 |
| cohort | Outcome | Predictor | estimate | Z-stat | P-value |
| ALSPAC | missingness | ODD/CD |  |  |  |
|  | age 10 | age 7 | 0.11 | 2.971 | 0.003 |
|  | age 13 | age 10 | 0.28 | 7.23 | < 0.001 |
|  | age 15 | age 13 | 0.16 | 5.56 | < 0.001 |
| ALSPAC | missingness | ADHD |  |  |  |
|  | age 10 | age 7 | 0.067 | 2.365 | 0.018 |
|  | age 13 | age 10 | 0.193 | 6.876 | < 0.001 |
|  | age 15 | age 13 | 0.11 | 4.152 | < 0.001 |
| ALSPAC | missingness | Depression |  |  |  |
|  | age 10 | age 7 | 0.026 | 0.637 | 0.0525 |
|  | age 13 | age 10 | 0.093 | 2.55 | 0.011 |
|  | age 15 | age 13 | 0.018 | 0.596 | 0.55 |
| ALSPAC | missingness | Anxiety |  |  |  |
|  | age 10 | age 7 | -0.05331 | -2.179 | 0.029 |
|  | age 13 | age 10 | 0.025 | 1.01 | 0.312 |
|  | age 15 | age 13 | -0.011 | -0.0518 | 0.604 |

The polygenic risk score based on the polygenic prior set to 1 was used. Results did not vary strongly based on the prior.

## Supplementary Note 1

This note accompanies the manuscript entitled: “Genetic overlap between schizophrenia and developmental psychopathology: a longitudinal analysis of common childhood disorders between age 7 and 15”. All data described here and analyses presented here serve to support the conclusions of the manuscript as published.

**Phenotypic descriptive statistics**

**Supplementary Table 2** presents the mean scores on the DSM-IV based scales of anxiety, depression, ADHD, ODD/CD for males and females in the NTR at different ages (left) as well as the percentages of male and female ALSPAC participants with these diagnoses, defined as a score of 4 or 5 on the DAWBA (right). (Note that in the analyses, the 6-category DAWBA band was used as outcome variable since this is a more informative measure than the dichotomous DAWBA diagnosis)

**Genotyping and genotype quality control:**

The NTR participants were genotyped on Affymetrix 6.0, Affymetrix-perlegen 5.0, Illumina 660 and Omni express (1M) platforms. Array specific calls and cleaning were performed before data from different platforms were combined. Data from different platforms were strand aligned, SNPs with a minor allele frequency below 1%, a HWE p-value < 1*10^−5^ and with a genotype missingness rate > 10% or call rate < 95% were removed. Individuals with an excessive or low heterozygosity were removed (F > .10 or F < .10). After QC, genotypes were imputed to a common set of SNPs based on the goNL reference set.^1^ SNPs were imputed that were not directly measured on eachplatform. Samples were excluded when reported gender did not match biological gender or when individuals were of non-European ancestry based on principle component analysis.^2^ In the NTR, sex, call rate, F (inbreeding coefficient), five principle components based on global ancestry and five principal components correcting for local ancestry differences within the Netherlands were included as covariates in all analyses.^2^

In ALSPAC, children were genotyped on the Illumina HumanHap550 quad chip genotyping platforms. The raw genome-wide data were subjected to standard quality control methods. Individuals were excluded on the basis of gender mismatches, minimal or excessive heterozygosity, disproportionate levels of individual missingness (>3%), and insufficient sample replication (IBD < 0.8). Population stratification was assessed by multidimensional scaling analysis, and compared with Hapmap II (release 22); all individuals of non-European ancestry were removed. SNPs with a minor allele frequency of < 1%, a call rate of < 95%, or evidence for violations of Hardy-Weinberg equilibrium (p < 5E^−7^) were removed. Cryptic relatedness was measured as proportion of identity by descent (IBD > 0.1). Related subjects that passed all other quality control thresholds were retained during subsequent phasing and imputation. Imputation of the target data was performed using Impute V2.2.2^3^ against the 1000 genomes phase 1 version 3 reference panel, using all 2186 reference haplotypes (including non-Europeans)^4^. As the ALSPAC sample, after QC, is assumed to be genetically homogeneous with respect to ancestry and local ancestry differences, no principal components were added to as covariates, sex was included as covariate in all analyses.

**The correction for the presence of overlapping subjects at the different ages of measurement, and the correlation between the polygenic predictors**.

In the meta regression analysis performed we have a set of 6 predictors X, and 32 outcomes Y. We perform a series of 192 univariate regressions:

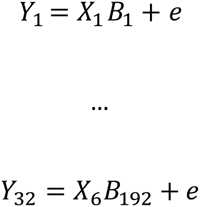

We construct an approximate error correlation matrix (i.e. the correlation between the regression parameters B) for a series of univariate regressions of p equal to:

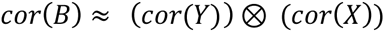

We specify the error *covariance* matrix as: *se* * *cor*(*B*) * *se*. Where *se* was a 192 × 192 diagonal matrix with the standard associated with each of the parameters Bon the diagonal. The errors were assumed to be independent between cohorts and therefore correlations between cohorts were set to zero in matrix *cor*(*B*). Based on the specified error correlation matrix, we performed the meta-analysis and meta-regression of the beta’s obtained from the univariate regression analyses. To test the proposed error correlation matrix accurately accounts for the dependence induced by correlated predictors and outcomes, we performed type 1 error simulations.

We simulated 3 traits (Y) (correlations between .3 and .5) and 3 polygenic scores (X) (correlations between .9 and .8) for 100 subjects. In each simulation there is no true association between PRS and traits. We regress each trait Y on each polygenic score X, and meta analyze the 9 test statistics obtained from these regressions, correcting for the dependence between traits and risks cores as outlined above. Given a small sample in the univariate regressions (N=100) the following slightly liberal type 1 error rates were observed. The liberal type-1 erro is likely induced by the fact that the test statistic obtained in each meta analysis follows at-distribution and not a normal distribution.

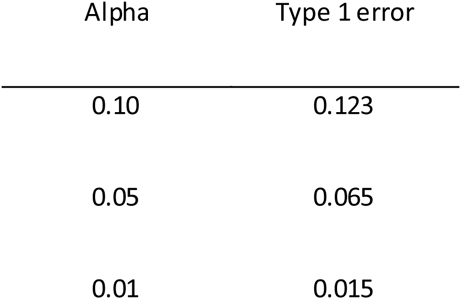

Simulating data given a larger sample of 1000 subjects in the initial univariate regressions the following, accurate, type 1 error rates were observed:

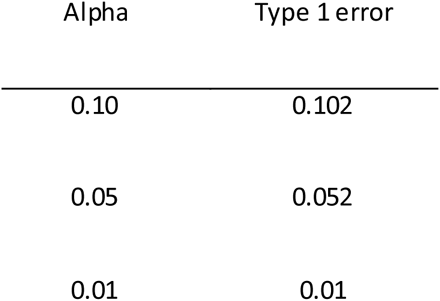

In our study, the sample size for the individual univariate regressions to be meta-analyzed ranged between 1200 and 6000 thus we were satisfied with the results of the type-1 error simulations.

A different limitation was that the error covariance as specified here assumed total sample overlap, and the absence of any covariates. Note however we do include covariates to control for population stratification and mean differences between male and female participants and these effects are assumed to be sufficiently small to allow our approximation to be valid. Strong covariate effects and substantial dropout would likely reduce power to detect an overall or age effect, and possibly increase type 1 error in some situation. As a form of sensitivity analysis the off diagonal elements of the phenotypic correlation matrices in ALSPAC and NTR were shrunk by 50% or 33% and increased by up to 10% to simulate the effect of less than total sample overlap or the effect of covariates changing the error covariance matrix. The substantive conclusions remained virtually unchanged, model 3 as described in the main text, had the best model fit when the off-diagonal elements in the phenotypic covariance matrix were reduced 50 or 33%, model 4 performed best when the phenotypic covariance was increased 10%. Parameter estimates and test statistics in model 3, fitted on the increased or decreased error covariance matrix were virtually unchanged. The sensitivity analyses revealed the effects of misspecification of the error covariance matrix did in all likelihood not influence the conclusions.

**Parametric resampling of the data to account for the influence of sampling fluctuation**

To quantify the influence of sampling fluctuation on our model selection we resampled the input for the meta-regression from a multivariate normal distribution with means equal to the observed regression coefficients in the univariate PRS analyses, and covariance equal to the above specified error covariance matrix. Unlike non-parametric bootstrapping this technique does make assumptions about the asymptotic distributions of test statistics. However parametric resampling does allow for quantification of sampling variance in the model selection procedure.

**Mixed effects meta regression to account for residual heterogeneity**

The best fitting fixed effects meta analysis model (model 3; **Table 1**) revealed a moderate amount of residual variation not accounted for by the meta regression model (Qe =226,49, df=181, p = 0.0122). We therefore performed 2 additional random effects meta analyses. Our First random effect model allowed for (correlated) random effects for each observed effect size, where the correlations between the random effect were assumed equal to the error covariance. The first mixed effects model significantly improves the model fit (LRT=8.81, df = 1, p = 0.0009). The fixed effects age, ODDCD, and agexADHD are all significant (p <0.05) in this mixed effects model (as they were in the mixed effects meta-regression model) the effect agexODDCD no longer reaches significance (p = 0.0681). The omnibus test of all meta-regression parameters combined also remained significant (QM = 37.0610, df = 10, p < 0.0001). The second mixed effects model includes a random intercept (i.e. In this model, the only dependence between effect sizes is introduce by the meta regressors or the error covariance. This second random effects model also significantly improves model fit over the fixed effects model (LRT = 14.7982, df=1, p < 0.0001). The second mixed effects model reveals a significant age effect and agexADHD effect (p< 0.05), but no significant ODDCD and agexODDCD effects. The overall test of moderators remains significant (QM= 30.0032, df=10, p = 0.0009. In general both mixed effects models retain the main conclusions of the fixed effects model (i.e an increasing association between schizophrenia PRS and childhood psychopathology with age, and some differences between the disorders in their relationship with schizophrenia).

**Estimating genetic correlations based on polygenic risk score results**

The univariate polygenic risk analyses results were obtained from either an ordered logistic regression (ALSPAC) or general estimation equations (GEE in NTR). For explanatory purpose we consider an OLS regression:

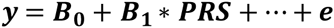
where the trait (y) and the PRS are scaled to unit variance and centered. The regression contains a number of other covariates (such as sex and principal components). Assuming the effects of the principal components and sex on the phenotypes are small to negligible, the square of B1 (**B**_1_^2^) is equal to the variance explained in the phenotype by the PRS. We further assume that the squared predicted outcome of the multivariate meta-analyses correspond to R^2^. Given these assumptions we used the previously derived relationship between R^2^ and genetic correlation^5^ to approximate the genetic correlations between childhood psychopathology and schizophrenia:

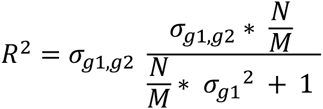

The inverse relationship equals:

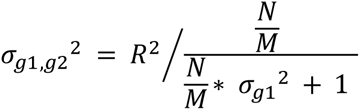

From which we can obtain:

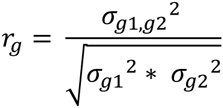
where N equals sample size in the discovery sample, M equals the independent number of genetic effects in the set of SNPs, σ_*g*1,*g*2_^2^ is the genetic covariance between target and discovery trait and σ_*g*1_^2^ equals the genetic variance explained by all measured markers in the target trait. As the discovery sample here is an ascertained case control sample (34241 cases and 45604 controls) we substitute the effective N using the effective sample size formula proposed by Wilier et al.^6^

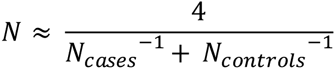

Note that N is approximate and R^2^ is estimated directly in the PRS analyses. Therefore, we needed to assume values for M and σ_*g*1_ ^2^ Any uncertainty in these values will not be reflected in the confidence bounds around the genetic covariance. We assume M to equal 200.000, to explore the influence of the uncertainty in σ_*g*1_ ^2^ on the estimate of r_g_ we computed the genetic correlation assuming the heritability explained by all SNPs included in the score for childhood psychopathology is 0.15 (**Figure 3**), .20 (**Supplementary Figure 1**) or .10 (**Supplementary Figure 2**). Note that we do not account for differences in the variance explained by the SNPs for the different psychopathologies at the different ages. We further assumed that the equations remain valid for estimates of B obtained from GEE (to correct for the presence of related samples) or ordered logistic regression (to correct for the fact that the ALSPAC phenotype was an ordered categorical variable).

